# Agri-environment nectar chemistry suppresses parasite social epidemiology in an important pollinator

**DOI:** 10.1101/2021.01.30.428928

**Authors:** Arran J. Folly, Hauke Koch, Iain W. Farrell, Philip C. Stevenson, Mark J.F. Brown

## Abstract

Emergent infectious diseases are a principal driver of biodiversity loss globally. The population and range declines of a suite of North American bumblebees, a group of important pollinators, have been linked to emergent infection with the microsporidian *Nosema bombi*. Previous work has shown that phytochemicals in pollen and nectar can negatively impact parasites in individual bumblebees, but how this relates to social epidemiology and by extension whether plants can be effectively used as disease management strategies remains unexplored. Here we show that caffeine, identified in the nectar of Sainfoin, a constituent of agri-environment schemes, significantly reduced *N. bombi* infection intensity in individual bumblebees and, for the first time, that such effects impact social epidemiology, with colonies reared from wild caught queens having lower prevalence and intensity of infection. Furthermore, infection prevalence was lower in foraging bumblebees from these colonies, suggesting a likely reduction in population-level transmission. Our results demonstrate that phytochemicals can impact pollinator disease epidemiology and that planting strategies, which increase floral abundance to support biodiversity could be co-opted as disease management strategies.

---

Two prominent drivers of biodiversity loss are emerging infectious diseases (EID) [1 – 7] and the reduction of natural habitat [8 – 13], often as a direct consequence of intensive agriculture [14 – 16]. One important approach to mitigate the negative impact of agriculture on biodiversity has been the development of agri-environment schemes (AES) [17,18] and conservation reserve programmes (CRP) [19]. In both AES and CRP, prescriptions are set out that increase floral abundance and diversity to conserve and enhance broader biodiversity, and the ecosystem services, such as pollination, that it supplies [17 – 19]. Such approaches have been shown to increase the abundance, diversity and persistence of beneficial organisms, such as pollinators, in agricultural environments [20 – 23]. However, pollinators are also threatened by EIDs [24, 25], and AES prescriptions may indirectly act as hubs that amplify disease in these populations [25 – 27]. Floral rewards, the main attractant for pollinators, contain secondary metabolites [28 – 30]. These phytochemicals can have antimicrobial properties [31] and may therefore have positive effects on pollinator disease by controlling parasites and pathogens [32 – 36]. Consequently, strategies that increase floral abundance and diversity, if designed to include floral mixes that incorporate high nutritional and medicinal value, may improve pollinator health and therefore could be co-opted to manage wildlife diseases.

Bumblebees are key global crop pollinators [37 – 39]. However, these charismatic insects are threatened by EIDs [40, 41]. More specifically, disease spillover has been identified between the European honeybee (*Apis mellifera*) and UK bumblebees [25, 42] and the global movement of commercial bumblebees for crop pollination has introduced novel pathogens to naive bumblebee communities in both North and South America [24, 43]. One such emergent pathogen is the microsporidian *Nosema bombi* [44], which has been implicated in the population and range declines recorded in a suite of North American bumblebees [24, 45]. Bumblebees are unable to recognise and remove *N. bombi* infected brood [46], suggesting that social immunity [47] is insufficient to adequately prevent disease transmission within a colony. Given that animals consume naturally occurring, bioactive compounds in their diets [48 – 50], inspiration for novel medications to combat parasitic infections may be drawn from an animal’s pre-existing diet [51 – 53], as has been shown to be the case for individual bumblebees [34 – 36]. However, such studies have yet to address the social epidemiology of pollinator parasites, which is key to their success in highly social insects [54].

Here we screen flowers used in UK-based AES, targeted at bumblebees, for phytochemicals in both the pollen and nectar. Caffeine was identified in the nectar of sainfoin (*Onobrychis viciifolia*), a major global crop [55]. We then designed a range of experiments to determine if consumption of caffeine can impact *N. bombi* infection in individual bumblebee workers, both prophylactically and therapeutically. We then used colonies reared from wild bumblebee queens to investigate whether caffeine can impact social, and by extension environmental epidemiology. Given that phytochemicals impact disease in individual bumblebees, we predict that if caffeine has a negative impact on *N. bombi* in individuals it will also impact social epidemiology.

## Results

### Identification and analysis of phytochemicals from AES plants

Nine species of flowers were identified from AES seed mixes prescribed by Natural England (2017) as potential bumblebee forage (supplementary table 1). Forty-one and twenty-seven phytochemicals (inclusive of isomers) were identified respectively from pollen and nectar sampled from the nine AES species, using LC-MS (supplementary table 2 & supplementary table 3 respectively). The alkaloid caffeine was identified in the nectar of sainfoin (*Onobrychis viciifolia*) (concentration range 0.35-200μM) using a retention time (rt) of 6.11 and a mass-to-charge ratio (*m/z*) of 195.09. For all experimental procedures we selected 200μM of caffeine to investigate its impact on *N. bombi*. While this was the highest concentration recorded in our analyses, from a limited number of sampling points and locations, it was representative of natural caffeine concentrations reported elsewhere in floral nectar [56, 57].

### Caffeine negatively impacts *N. bombi* infection in *B. terrestris* workers

In the prophylactic bioassay, 82 workers successfully eclosed, of which 36 had *N. bombi* infections (control = 22, caffeine = 14), resulting in an overall infection success of 44%. There were no significant differences between the infection prevalence in our control or experimental groups (χ^2^ = 1.433, *P* = 0.231), indicating that caffeine had not affected parasite prevalence. In contrast, caffeine did have a significant prophylactic effect by reducing *N. bombi* infection intensity in eclosed workers (LMM, *F*_1,28_ = 15.33, *P* < 0.001; Figure 1a). The covariates thorax width (LMM, *F*_1,28_ = 3.75, *P* = 0.06), faeces volume (LMM, *F*_1,28_ = 0.65, *P* = 0.4) and the random effect colony (*P* = 0.27) all had no significant effect on *N. bombi* infection intensity.

**Figure 1.**
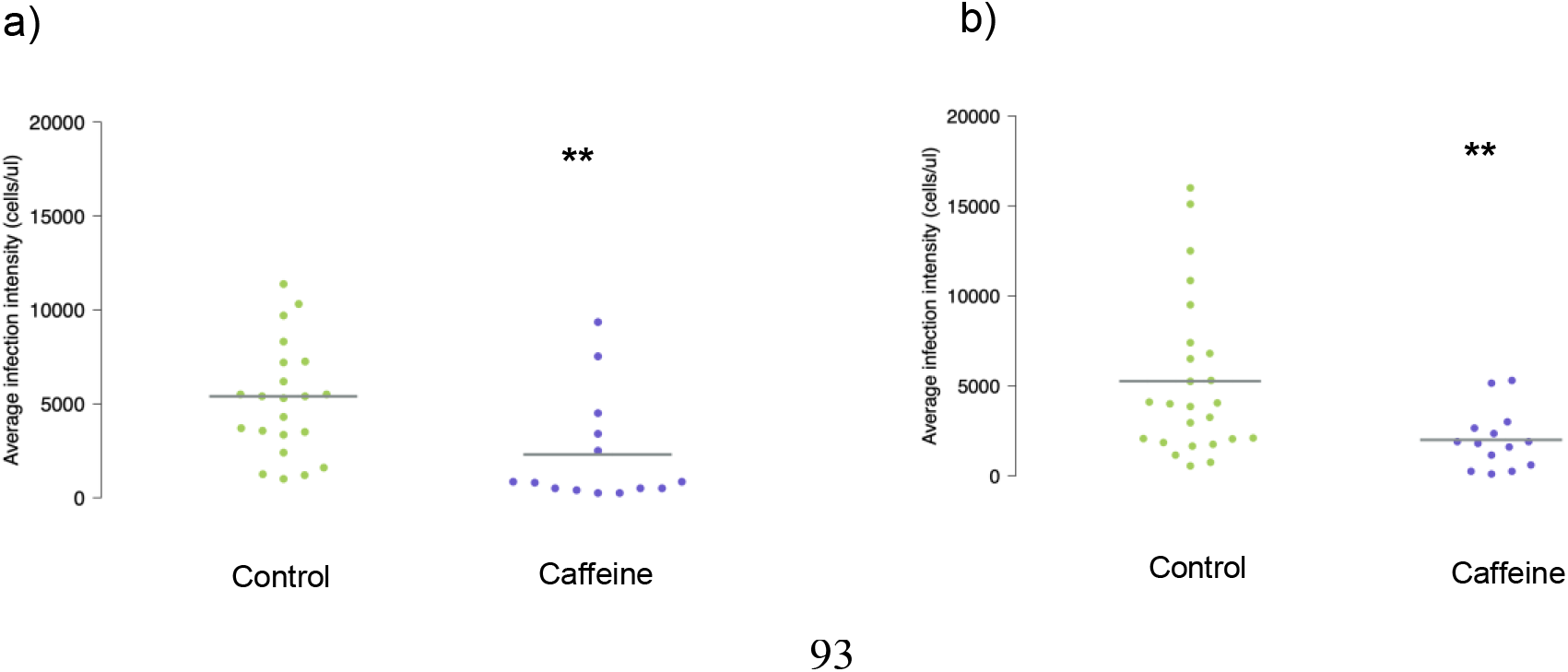
Beeswarm plots of *N. bombi* infection intensities in adult *B. terrestris* workers that had been fed caffeine prophylactically (a) and therapeutically (b). The sample mean has been marked with a grey bar and statistical differences have been marked with a double asterisk. Caffeine was found to have both a significant prophylactic and therapeutic effect on *N. bombi* infection intensity in *B. terrestris* workers.

In the therapeutic bioassay, 101 workers successfully eclosed, of which 39 had *N. bombi* infections (control = 25, caffeine = 14), resulting in an overall infection success of 39%. Again, there was no significant difference between the infection prevalence in our control or experimental groups (caffeine χ^2^ = 2.384, *P* = 0.123). However, as with our prophylactic bioassay, caffeine had a significant therapeutic effect by reducing *N. bombi* infection intensity in eclosed workers (LMM, *F*_1,31_ = 4.97, *P* = 0.032; Figure 1b). The covariates thorax width (LMM, *F*_1,31_ = 2.49, *P* = 0.123), faeces volume (LMM, *F*_1,31_ = 1.87, *P* = 0.181), and the random effect colony (*P* = 0.193) all had no significant effect on *N. bombi* infection intensity.

### Caffeine significantly impacts the epidemiology of *N. bombi* in *B. terrestris* colonies

Caffeine treatment significantly reduced the prevalence of *N. bombi* in colonies (LMM, *F*_1,851_ = 18.04, *P* < 0.001; Figure 2). Prevalence varied across colonies (*P* = 0.001) but the covariates brood cohort (LMM, *F*_1,851_ = 3.36, *P* = 0.06), number of inoculated larvae (LMM, *F*_1,851_ = 0.02, *P* = 0.90), number of days since inoculation (LMM, *F*_1,851_ = 0.23, *P* = 0.63) and the random effect forager (*P* = 0.06) all had no significant effect on *N. bombi* infection prevalence. Caffeine had a similar negative impact on *N. bombi* infection intensity (LMM, *F*_1,851_ = 34.8, *P* < 0.001; Figure 3). In addition, there was a significant negative interaction between cohort and treatment on *N. bombi* infection intensity (*F*_1,851_ = 14.2, *P* < 0.001), meaning that the younger the bumblebee, the greater the impact caffeine treatment had on reducing *N. bombi* infection intensity (Figure 4). In contrast, the covariates number of inoculated larvae (LMM, *F*_1,851_ = 0.24, *P* = 0.6), number of days since inoculation (LMM, *F*_1,851_ = 0.001, *P* = 0.9) and the random effects forager (*P* = 0.06) and colony (*P* = 0.51) all had no significant effect on *N. bombi* infection intensity (Figure 3). Finally, the covariates treatment (LMM, *F*_1,205_ = 0.937, *P* = 0.02) and cohort (LMM, *F*_1,205_ = 2.294, *P* < 0.001) were found to have a significant impact on forager infection prevalence. In essence foragers, especially younger foragers, from our caffeine treated colonies were less likely to have *N. bombi* infections. In contrast, the infection intensity of infected foragers was not impacted by treatment (LMM, *F*_1,205_ = 1.509, *P* = 0.14), but interestingly cohort did have a significant negative impact on forager infection intensity (LMM, *F*_1,205_ = 3.061, *P* <0.001), showing that infected younger foragers had lower infection intensities (Figure 5). In both forager models faeces volume (Prevalence LMM, *F*_1,205_ = −1.505, *P* = 0.13, Infection intensity LMM, *F*_1,205_ = 0.342, *P* = 0.7), number of larvae inoculated (Prevalence LMM, *F*_1,205_ = −0.242, *P* = 0.8, Infection intensity LMM, *F*_1,205_ = 01.323, *P* = 0.19) and colony (Prevalence LMM *P* > 0.05, Infection intensity LMM *P* > 0.05) had no impact on infection prevalence or intensity.

**Figure 2.**
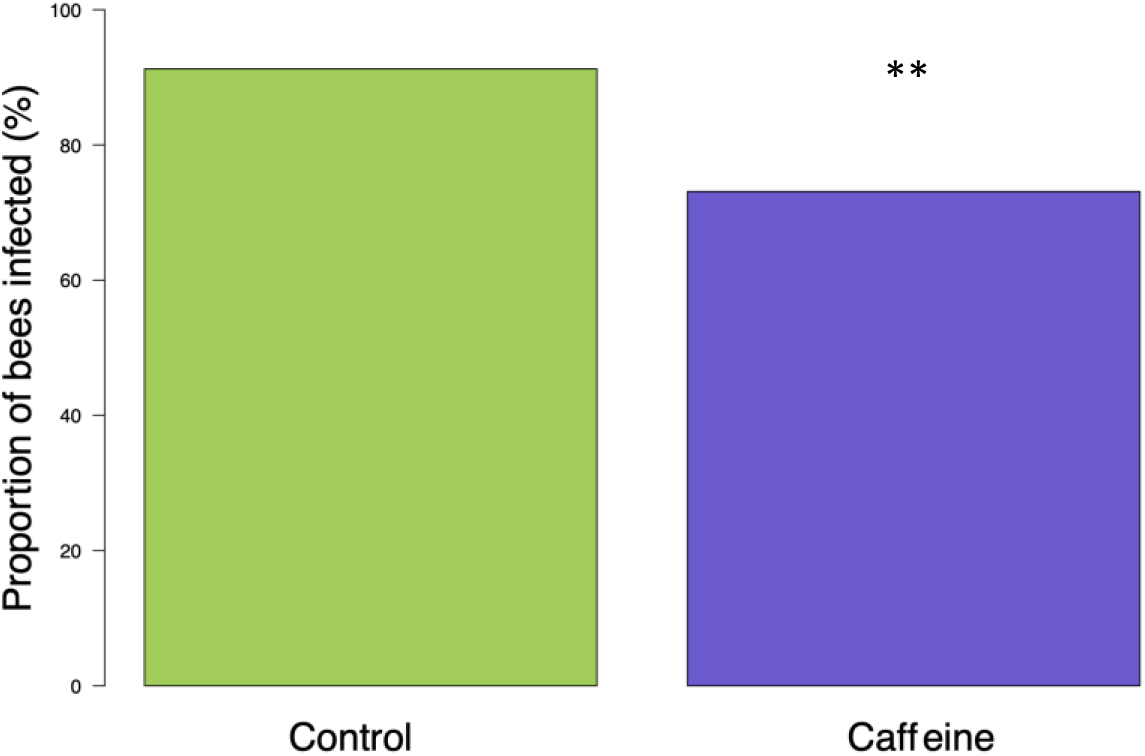
*Nosema bombi* infection prevalence in adult bumblebees (*B. terrestris*) for both control (n = 428) and caffeine (n = 272) treatments. Treatment with caffeine significantly reduced *N. bombi* infection prevalence.

**Figure 3.**
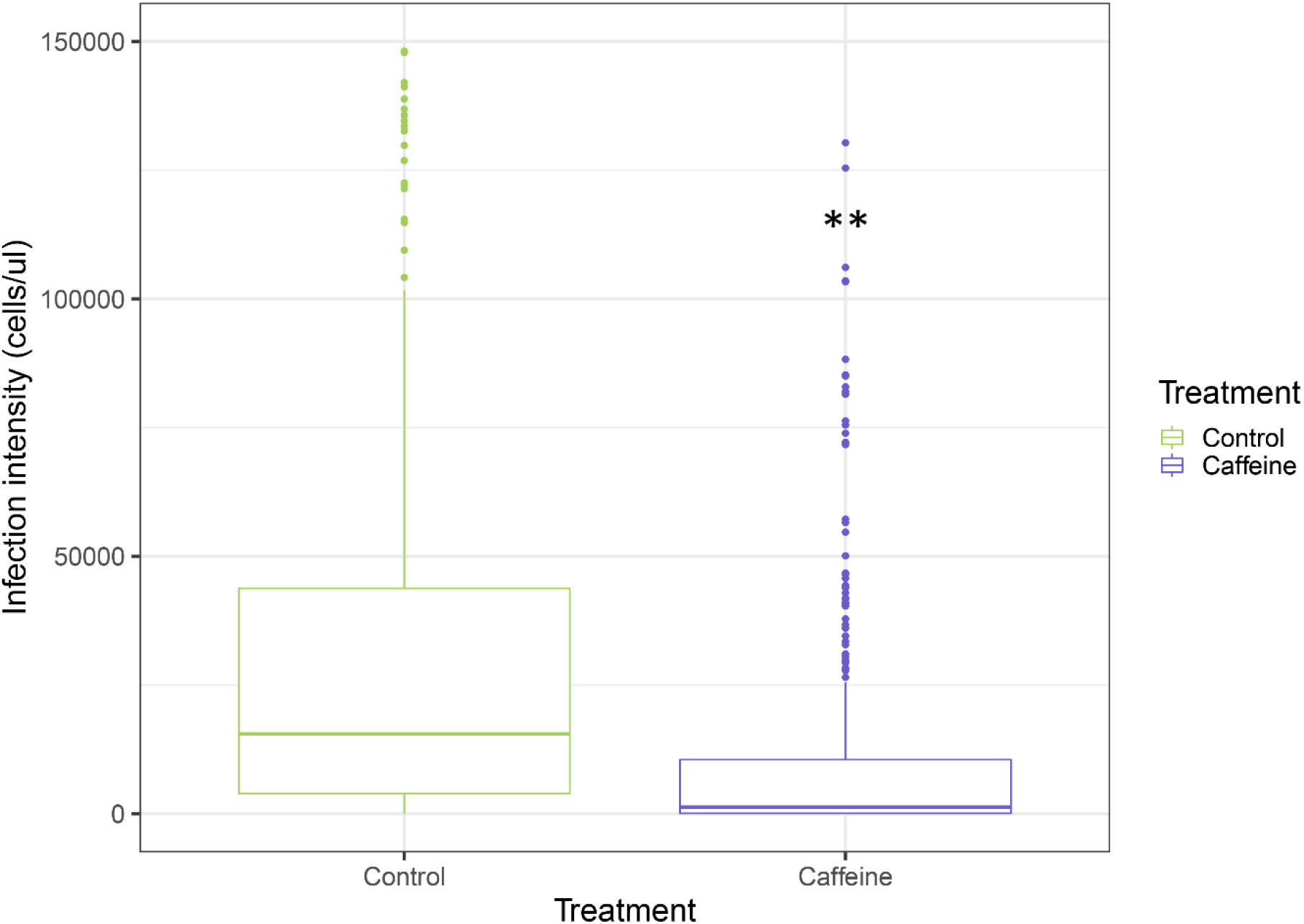
Treatment with caffeine significantly reduced *N. bombi* infection intensity. Box plot of *Nosema bombi* infection intensity in adult bumblebees (*B. terrestris*) for both control (n = 438) and caffeine (n = 272) treatments. Significant differences between treatments are shown with a double asterisk.

**Figure 4.**
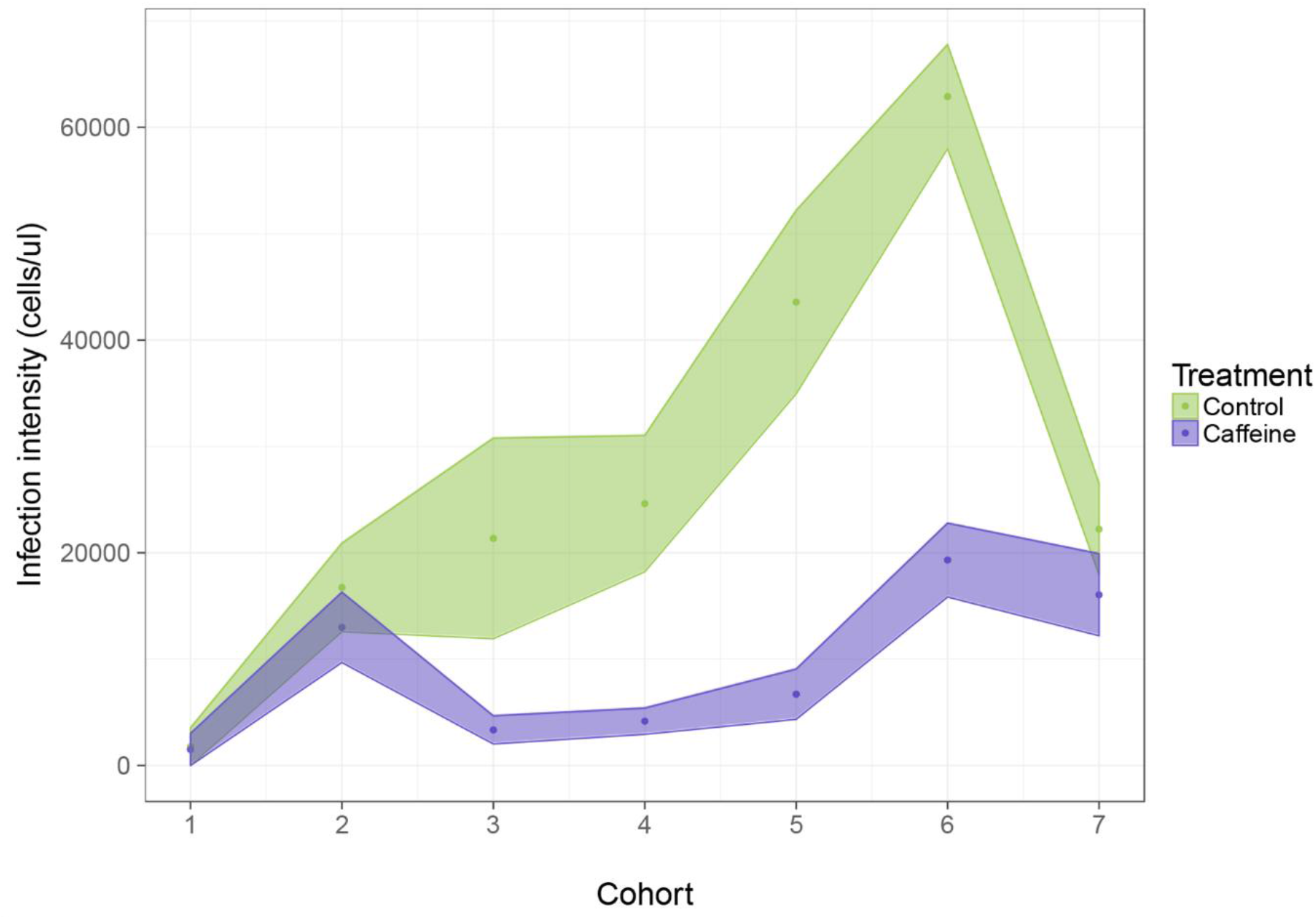
*Nosema bombi* infection intensity across *B. terrestris* brood cohorts. Each brood cohort is a representation of a colonies reproductive output over two weeks, with the first cohort representing the first batch of brood produced. Hence, at sampling, later cohorts contained younger bumblebees. Cohort and treatment significantly impacted *N. bombi* infection intensity in adult bumblebees. Caffeine feeding reduced *N. bombi* infection intensity and earlier cohorts across both treatments had lower infection intensities. Shaded areas represent mean ±SEM.

**Figure 5.**
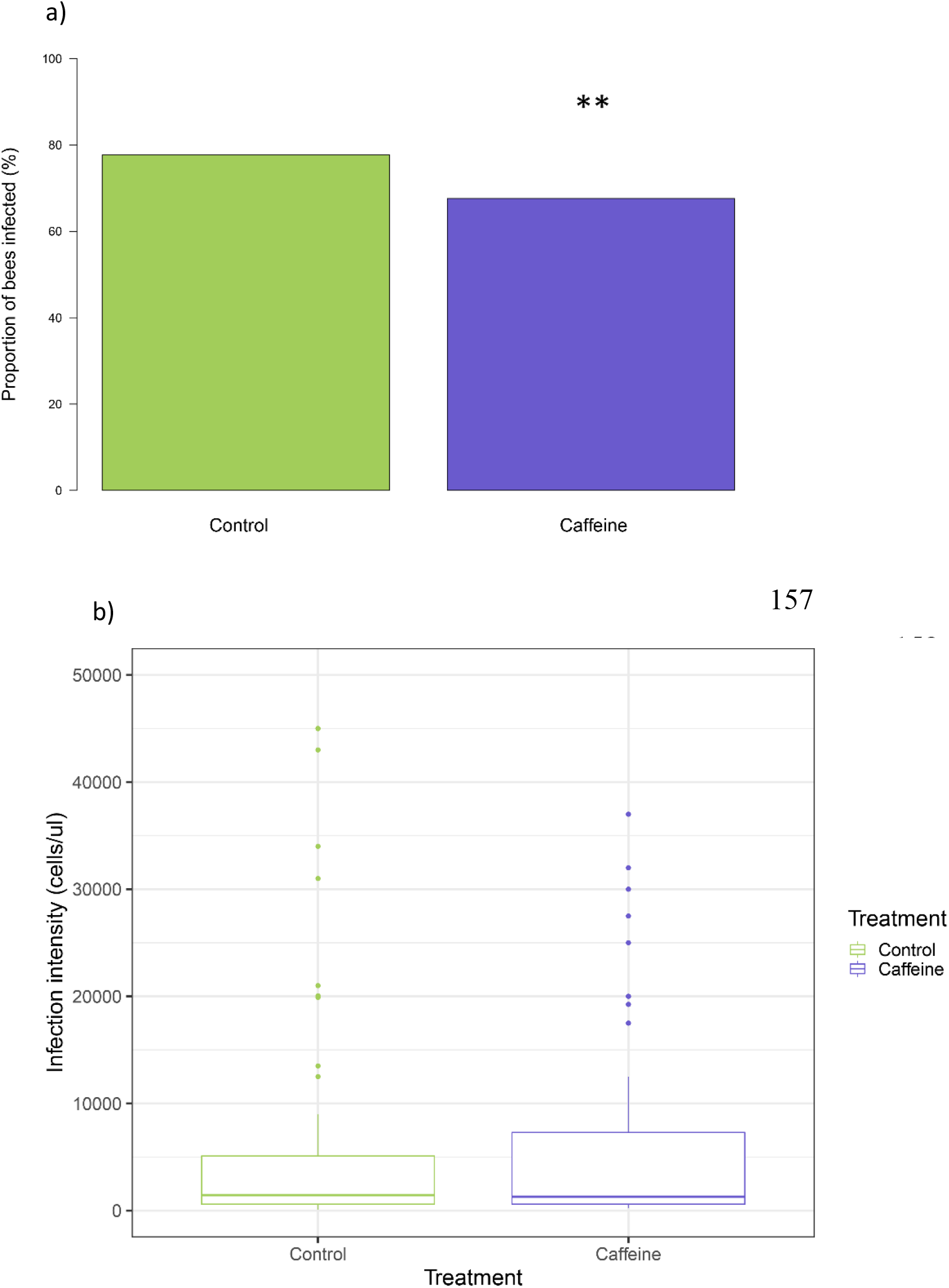
*Nosema bombi* infection prevalence (a) and infection intensity (b) for foraging bumblebees sampled during the experiment. There were significantly fewer bees that had *N. bombi* infections in the caffeine treatment when compared to the control group. In contrast, there was no significant difference in infection intensities between the caffeine and control groups.

### No impact of caffeine on bumblebee colony growth and reproduction

The covariates treatment (LMM, *F*_1,18_ = 2.45, *P* = 0.13), number of brood inoculated (LMM, *F*_1,18_ = 0.77, *P* = 0.3) and number of days since inoculation (LMM, *F*_1,18_ = 0.004, *P* = 0.4) had no effect on the number of workers produced in *B. terrestris* colonies. Similarly, the covariates treatment (LMM, *F*_1,18_ = 0.96, *P* = 0.3), population size (LMM, *F*_1,18_ = 2.13, *P* = 0.17), number of brood inoculated (LMM, *F*_1,18_ = 0.04, *P* = 0.85), days since inoculation (LMM, *F*_1,18_ = 0.032, *P* = 0.25) and the random effect colony (P = 0.82) had no effect on the production of sexual castes.

## Discussion

Here we show that phytochemicals which may be encountered by bumblebees in AES landscapes can negatively impact both the individual and social epidemiology of a key bumblebee pathogen, *N. bombi*. Specifically, consumption of caffeine reduced the overall infection prevalence and intensity of *N. bombi* infections in *B. terrestris* colonies reared from wild caught queens, while simultaneously having no negative impact on colony development. Consequently, our results indicate that schemes that increase floral abundance and diversity, such as AES and CRP may have unexplored benefits as they have the potential to be co-opted as disease management tools to further improve pollinator health in the field.

We identified caffeine in the nectar of sainfoin (*Onobrychis viciifolia*), a component of UK AES, and showed that it negatively impacts the infection success and epidemiology of the microsporidian *N. bombi* in wild caught and reared *B. terrestris* colonies. While our assessments of caffeine bioactivity were at the highest concentration likely to be encountered in wild *O. viciifolia*, this is still within naturally occurring nectar concentrations reported from other caffeine producing plants [56]. In addition, caffeine is a common nectar constituent [57]. Consequently, our experimental paradigm represents an ecologically relevant interaction between pollinators, their pathogens and forage plants. However, we would note that further work is required to assess the variation across different cultivars and wild species of caffeine producing plants [58].

Emergent infection with *N. bombi* has been implicated as a principal driver of contemporary bumblebee declines across North America [24, 45]. Our results show that caffeine reduced the overall infection intensity of *N. bombi in vivo*, in individual bumblebees that were treated as larvae, in both prophylactic and therapeutic treatments. Both of these treatment methods are recognized in preventative medicine and are biologically realistic within the bumblebee-*Nosema* system. As infection by *N. bombi* may occur at any time across the lifecycle of bumblebee colonies, caffeine, and potentially other bioactive secondary chemicals [36], may interact either prophylactically or therapeutically to prevent or inhibit parasitaemia. Exposure to such compounds depends upon the duration of flowering and density of flowers, both of which can be high in AES schemes [20]. Caffeine is well known for its biological activity [59, 60], and its impacts on *N. bombi* infection may be through reducing spore germination and subsequent infection success. This is likely to be enhanced in bumblebee larvae, whose blind gut may lead to an increase in caffeine concentration during the exposure period, resulting in a greater impact on the parasite and contributing to the reduction of infection prevalence and intensity recorded in adult bumblebees following eclosure.

These impacts of caffeine on individual infections were magnified at the colony level, with treated colonies having significantly lower prevalence and intensity of infections. In control colonies, the prevalence and intensity of infections increased across the colony cycle including the sexual production phase, which has implications for environmental disease persistence [61]. In contrast, both prevalence and intensity remained at relatively lower levels in treated colonies. This will have reduced the force of intra-colonial infection, by lowering the number of infective spores in the social environment. It is also likely to have significant knock-on effects for transmission between colonies. As *N. bombi* is not species specific [62], the reduction in disease prevalence, recorded here, from caffeine consumption in foraging bumblebees may also reduce the likelihood of inter-specific disease transmission [63, 64], and thus further impact environmental disease epidemiology. Interestingly, the higher infection intensities we observed in younger cohorts in control colonies were sufficient to yield higher colony infection prevalence. We suggest that this early *N. bombi* bloom in young adults, which caffeine inhibited, either through larval or adult consumption, as both may impact transgenerational disease in bumblebees [36], is critical in maintaining high intracolonial disease prevalence. Consequently, our results suggest that beneficial forage plants, such as those that produce caffeine, if provided throughout the lifecycle of a bumblebee colony (either by overlapping or prolonged anthesis periods) may mitigate the impact of endemic and emergent disease. Moreover, it is important to note that the impact of the *N. bombi* bloom in young adult bumblebees on prevalence is likely to be greater in incipient colonies where there are fewer individuals. Consequently, younger bees will be involved in nest-based tasks including nursing behaviours [65], where they are likely to pass spores to developing larvae [66], thus contributing to disease amplification. As such, planting strategies that target incipient colonies may have a greater impact on pollinator parasite epidemiology. The emergence of *N. bombi* in North America has been implicated as a driver of indigenous bumblebee declines [24]. Sainfoin is grown in North America and caffeine is present in the nectar of other plants found throughout North America [57]. Consequently, we recommend our results be studied in the context of North American bumblebee declines in response to EIDs. Our findings suggest that the incidence and impact of *N. bombi* can be mitigated with adaptive planting strategies, which include forage plants containing caffeine. Alongside the management of endemic disease, this approach may be implemented to provide a mechanism with which to minimize the spread of emergent disease to novel areas.

Infection with *N. bombi* dramatically impacts bumblebee colony health and fitness [67, 68] and this may have implications for population persistence [24]. Phytochemicals can negatively impact bumblebee fitness [69]. In contrast, our results show that chronic caffeine consumption did not have a negative effect on the health or fitness of our experimental colonies, indeed caffeine treated colonies had a trend for longer persistence and the production of more individuals of sexual castes in contrast to what would typically be expected in *N. bombi* infected colonies [68]. Consequently, at the colony level, caffeine is not having a detrimental effect on brood development or on queen fecundity. Caffeine has been reported in thirteen orders of plants [57]. Consequently, our findings on the impact of caffeine on bumblebee parasite epidemiology may be relevant in other landscapes, globally, where AES and sainfoin are not present. In addition, caffeine has been shown to enhance a pollinator’s memory of reward [56], resulting in increased visitation rates by altering the foraging behaviour of bees. Our results suggest that such manipulation of pollinator behaviour by caffeine, resulting in repeated flower visitations [70], may also indirectly benefit foraging bumblebees by reducing the incidence and distribution of a key parasite, *N. bombi*, within a given environment. However, it should be noted that under field conditions caffeine consumption may lead to suboptimal foraging strategies [71] which may have a negative impact on bumblebee colony fitness through nutritional deficiencies.

The current guidelines and floral recommendations for AES have been developed with the scientific community [72, 73] and AES have been shown to increase bumblebee species richness [20, 21] and more recently bumblebee reproductive fitness [23]. By integrating epidemiological studies, such as the results presented here, into AES, and similar schemes across the globe, such as CRP, their management could be further adapted to include species that have beneficial floral chemistry, which may provide indirect fitness benefits to pollinators through disease management.

## Methodology

### Identification and analysis of phytochemicals from AES plants

Nine species of flowers were identified from AES seed mixes prescribed by Natural England (2017) as potential bumblebee forage (supplementary table 1). Fresh pollen and nectar were collected from Langridge, UK (ST741694), Salisbury Plains, UK (SU069440), Royal Botanic Gardens, Kew (RBG Kew), UK (TQ184769), and Egham, UK (TQ010709) between May 2016 and July 2017. This collection period enabled repeat sampling of species and also allowed for sampling of any species that may have already undergone anthesis in 2016. Flowers were initially covered using a muslin cloth and a cable tie for up to 24 hours. This ensured maximum harvest of both pollen and nectar [74]. Nectar was collected using a 2μl glass micro-capillary tube [75] and stored in a 1.5 ml UV protective Eppendorf tube. Pollen was removed by positioning a stamen over a separate 1.5 ml UV protective Eppendorf tube and rocking the target flower backwards and forwards. Both pollen and nectar samples were kept in a chilled container in the field prior to being stored in a −20°C freezer for subsequent analysis. For each species of plant, at each index site, nectar and pollen samples were pooled separately, to ensure that any geographic variation in phytochemicals could be detected. In addition, pooling samples from multiple individuals of the same species increased the likelihood of detecting phytochemicals that are found at naturally low concentrations within both pollen and nectar.

Chemical analysis of floral rewards was undertaken using LC-MS. Each pollen sample was initially centrifuged at 6000 rpm for 1 minute before 100μl of 100% methanol (MeOH) was added. Pollen samples were then vortexed for 20 seconds prior to being stored at room temperature for 24 hours to ensure maximum extraction of phytochemicals into the methanol solvent. Nectar samples were centrifuged at 6000 rpm for 1 minute before 50μl of 100% MeOH was added and were available for immediate analysis after 20 seconds of vortexing [76].

Once prepared, as outlined above, a 50μl aliquot of each sample was placed into an individually labelled low absorption LC vial. LC-MS analysis was carried out using an LC elution program on a Thermo Scientific Ultimate 3000 LC and Velos Pro MS detector, with 5μl injection volume onto a Phenomenex Luna C18 (2) column (150 x 4.0mm id, 5μm particle size) held at 30°C. A gradient elution was employed consisting of a mobile phase of (A) MeOH (B) H_2_O and (C) 1% HCO_2_H in MeCN at a flow rate of 0.5mL min-1. Phytochemicals were then identified on the resulting ion chromatogram in Xcalibur 2.1.0 (2012) comparing retention times, molecular weight and UV absorption with data from an existing compounds library at RBG Kew. Phytochemicals that could not be identified from these parameters were subjected to High Resolution Electrospray Ionization Mass Spectrometry (HR-ESI-MS) using a Thermoscientific LTQ Orbitrap^™^ XL with a 5μl injection volume onto a Phenomenex Luna C18 (150mm x 3mm id, 3μm particle size) held at 30°C. A gradient elution was employed consisting of a mobile phase of (A) H_2_O (B) MeOH (C) 1% CH_2_O2 in CH_3_CN at a flow rate of 0.4mL min-1. HR-ESI-MS analysis facilitated identification of phytochemicals from fragmentation patterns of the mass spectra and accurate molecular formulae.

Forty-one and twenty-seven phytochemicals (inclusive of isomers) were identified respectively from pollen and nectar sampled from the nine AES species, using LC-MS (supplementary table 2 & supplementary table 3 respectively). The concentration of each phytochemical was calculated in parts per million (ppm) by comparison of UV peak areas with standard curves at relevant floral reward concentrations (10, 100, 500, 1000 ppm) and then converted to micromolar (μM) [29, 76, 77]. In the absence of a specific standard, compounds with a similar UV absorption, based on the same chromophore, were used as an approximation. The alkaloid caffeine was identified at a concentration of 200μM in the nectar of sainfoin (*Onobrychis viciifolia*) using a retention time (rt) of 6.11 and a mass-to-charge ratio (*m/z*) of 195.09 from a pooled sample taken in 2017. Caffeine has been reported previously to have biological activity against microbes [59], including reducing mortality in honeybees infected with Israeli acute paralysis virus [60]. Consequently, caffeine was an ideal compound to investigate the potential impact of AES phytochemicals on *N. bombi* epidemiology in *B. terrestris*. Identification of caffeine in nectar of a species in the Leguminosae was surprising since it has only been reported previously in Leguminosae once [78]. Therefore, to confirm the presence of caffeine in sainfoin, flower nectar was resampled as described above in 2019, using four independent investigators which resulted in three independently pooled samples of sainfoin nectar. Again, caffeine was identified in all three samples, although at lower concentrations than the 2017 sample (0.35, 8.3 and 16.4 uM), when run in isolation from the other samples using the same LC-MS method as before. Caffeine was absent from analysis immediately prior to and following each of our samples, indicating that the instrument was not contaminated with caffeine. However, caffeine has been reported in nectar of several plant species [57] from different plant families and may occur more widely than previously thought in plants. For all experimental procedures we selected 200μM of caffeine to investigate the impact on *N. bombi*. While this was the highest concentration found in our concentration range, from a small number of sampling points and locations, it is representative of naturally occurring caffeine concentrations, which is a widely available nectar compound [56, 57].

### *Nosema bombi* inoculum

To elucidate the effect of caffeine on *N. bombi*, we used a larval inoculation paradigm, as larvae are the most susceptible stage to infection [80]. A wild *B. terrestris* queen that was naturally infected with *N. bombi* was caught from Windsor Great Park, UK (SU992703) in 2016 [36]. The infected queen’s gut was isolated by dissection and homogenized in 0.01M NH_4_Cl. The resulting spore solution was centrifuged at 4°C for 10 minutes at 5000 rpm to isolate and purify the spore pellet as described in [79]. The pellet was resuspended in 0.01M NH_4_Cl and the *N. bombi* concentration was calculated using a Neubauer improved haemocytometer. To confirm the presence of *N. bombi*, and to ensure the microsporidium was not *N. ceranae*, as these two microsporidia can be easily confused under a light microscope, a sample of the inoculum was subjected to PCR using primers and the protocol outlined in [80]. The inoculum was then used to infect three commercially available *Bombus terrestris audax* colonies (Biobest, Belgium) from which further spores were harvested to be used to create inocula (described above). The resulting inocula were stored in a −80 freezer until they were required.

A larval *N. bombi* inoculant was prepared by combining inverted sugar water and pollen (3:1) to create an artificial worker feed as outlined in [66]. This was then combined in equal proportions (100μl: 100μl) with the *N. bombi* inoculum to create an experimental inoculant. Prior to any larval inoculation, workers from each micro-colony were removed for an hour. This resulted in larvae having no access to food and therefore experimental inoculation would be more likely to elicit a feeding response.

### Investigating the impact of caffeine on *N. bombi* infection on individual epidemiology

Eight *B. terrestris audax* colonies (hereafter referred to as donor colonies) (containing a queen, brood and a mean of 45 (± 6.5 S.E.) workers) were obtained from Biobest, Belgium. Colonies were kept in a dark room at 26°C and 50% humidity (red light was used for any colony manipulation). To ensure colonies were healthy and developing normally they were monitored for 7 days prior to use in any experimental procedures. This included randomly screening 10% of the workers every two days, from each colony, for common parasitic infections (*Apicystis bombi*, *C. bombi* and *N. bombi*) in faeces using a phase-contrast microscope set to ×400 magnification. No infections were identified in any of the eight donor colonies.

Micro-colonies were established by removing 8 patches of brood containing approximately 10 developing larvae (growth stage L2-3), from each of the eight donor colonies. Each of these patches of brood were placed in an individual 140×80×55mm acrylic box. These micro-colonies were each provisioned with *ad libitum* pollen and sugar water, and 3 workers from their original donor colony to provide brood care. All pollen used throughout the experiment was irradiated using UV light to remove any microbes. Prior to being entered into the experiment all brood-caring workers were individually marked using a coloured, numbered Opalith ^®^ tag and recorded so that they could be distinguished from newly eclosed workers.

To investigate if caffeine had any prophylactic properties, 16 micro-colonies (2 per donor colony) as described above were used. Prior to inoculation, control larvae were kept in their original micro-colonies (n=8) and provided *ad libitum* pollen and sugar water. However, in the experimental groups *ad libitum* pollen and sugar water containing caffeine (Sigma Aldrich CO750) at 200μM (n=8 micro-colonies) was provided for 7 days. Caffeine was added to sugar water using 4ml of 40% MeOH L^-1^ as a solvent, control colonies also had 4ml of 40% MeOH L^-1^ added. After seven days both experimental and control larvae at either instar 2 or 3 were artificially inoculated with 50,000 *N. bombi* spores in 4.3μl of inoculant (described above) using a 20μl pipette. The spore concentration in the inoculum is within ecologically relevant values for *N. bombi* spores in faeces and is infective to developing bumblebee brood (Rutrecht & Brown 2008, Folly *et al*. 2020). The larvae were left to consume the inoculum for 30 minutes, before being returned to their micro-colony.

In a simultaneous experiment, the therapeutic effect of caffeine was investigated. Here, 16 micro-colonies (2 per donor colony) as described above were used. In contrast to the prophylactic investigation, larvae in the therapeutic investigation were each inoculated with 50,000 *N. bombi* spores in 4.3μl of experimental inoculant, as described above, using a 20μl pipette, prior to experimental feeding. The larvae were again left for 30 minutes to consume the inoculum. The inoculated larvae were returned to their respective micro-colonies with brood-caring workers. Each control micro-colony (n=8) was provisioned with *ad libitum* pollen and sugar water. However, in the experimental groups *ad libitum* pollen and sugar water containing caffeine at 200μM (n=8 micro-colonies) was provided for 7 days. As before, phytochemicals were added to sugar water using 4ml of 40% MeOH L^-1^, control colonies also had 4ml of 40% MeOH L^-1^ of sugar water.

In both feeding trials larvae were allowed to develop naturally and pupate in their respective micro-colonies. Once eclosed, new workers were marked using a coloured, numbered Opalith ^®^ tag and individually quarantined for 3 days in an inverted plastic cup, which was modified with a hole that enabled a 15ml falcon tube to be inserted. The falcon tube contained control inverted sugar water diluted with double distilled H_2_O (50% w/w) that the newly eclosed workers could feed on. A quarantine period of three days was used to ensure that faecal samples were not heavily contaminated with pollen grains as these can obscure parasites and complicate parasite quantification. At the end of the quarantine period each worker had its thorax width measured (mm) as a proxy for bumblebee size, using a set of Mitutoyo™ digital calipers. In addition, each worker was isolated in a 25ml plastic vial where it provided a faecal sample, collected in a 10 μl glass capillary, the volume (μl) of which was recorded. Following this, each worker’s faecal sample was screened for *N. bombi* by microscopic examination using a phase-contrast microscope at ×400 magnification. If an infection was identified a Neubauer improved haemocytometer was used to quantify the parasite load. Workers were then sacrificed and stored in a labeled Eppendorf tube at −80°C.

### Investigating the impact of caffeine on the social epidemiology of *N. bombi*

Between February and April 2018, 250 wild, foraging *B. terrestris* queens were collected from Windsor Great Park, Surrey, UK (SU992703), using an entomological net. Each queen was isolated into an individual, labelled, 25ml perforated collection vial and chilled in an insulated bag, before being returned to the laboratory. All queens were then screened for common bumblebee endoparasites (described above, including *Sphaerularia bombi* as this infects queen bumblebees) via microscopic examination of faeces using a phase contrast microscope at ×400 magnification. Following this initial screen, apparently uninfected bumblebee queens were quarantined in an individual 127×67×50mm acrylic box, where they were fed *ad libitum* pollen and sugar water, in a dedicated bumblebee rearing room, which was kept at 26°C and 50% humidity. As before, all pollen used throughout the experiment was irradiated using UV light to remove any microbes. Seven days after the initial parasite screen each queen was rescreened for the common bumblebee endoparasites as above. This seven-day delay ensured that any incipient infections missed during the initial screen were identified once parasitaemia had increased to detectable levels [81]. All queens that were infected during either of the parasite screens (n = 15) were excluded from the experiment and released back to the original field collection site.

Uninfected queens (hereafter referred to as queens) were returned to their acrylic box in the dedicated rearing room (described above) and entered into the experiment. Unless otherwise stated, all colony manipulation was carried out under red light using sterile equipment and workbenches. All queens were provided with *ad libitum* sugar water from a gravity feeder and a pollen ball for nutrition and to encourage egg-laying. Pollen balls were created by combining pollen and sugar water (50% w/w) in a 20:1 ratio. This mixture was then subdivided into individual 15×15×15mm cubes. Sugar water and pollen balls were changed every seven days or if a queen had spoiled her resources, whichever came first. Queens were left to develop naturally, and reproductive output was monitored and recorded. Once the first set of workers had hatched, these were marked using numbered, coloured Opalith ^®^ tags (tag colour was unique to cohort, not individual or colony) and the remaining incipient brood was inoculated with *N. bombi* (as described above) and entered into one of two feeding regimes. Any queen that did not lay a clutch of eggs within eight weeks (n = 113) was excluded from the experiment and released back to the original field collection site. Of the 250 queens originally caught, 40 produced an incipient colony, of which 19 lasted for the duration of the experiment.

Following inoculation (described above), the original queen was allowed to re-associate with the brood for 5 minutes prior to the marked workers being returned. This step reduced the likelihood of the queen rejecting the brood later in the colony lifecycle (AJF personal observation). Following this, each incipient colony (including queen and any hatched workers) was placed into a 290×220×130mm plastic colony box. This habitation box was connected, via a plastic tube, to a separate 290×220×130mm plastic box that was used as a foraging arena. The foraging arena provided *ad libitum* pollen and experimental sugar water. Each control colony (n = 10 colonies) was provisioned with *ad libitum* pollen and sugar water for the duration of the colony lifecycle. In the experimental group *ad libitum* pollen and sugar water, containing caffeine at 200μM (n = 9 colonies) was provided for the duration of the colony life cycle. Caffeine was added to sugar water using 4ml of 40% MeOH as a solvent L^-1^, control colonies also had 4ml of 40% MeOH added L^-1^, of sugar water. Pollen and sugar water were changed every seven days, or once the colony had consumed all of the resources, whichever came first. To investigate if caffeine had any effect on *N. bombi* epidemiology, a range of colony specific measurements were taken throughout the colony lifecycle. Every two weeks all newly eclosed workers were individually marked using a coloured, high varnish paint, which was unique to that specific brood cohort and not the colony. Marking bumblebees in this way enabled investigation of colony demographics and infection prevalence and intensity. A period of two weeks was used to ensure that larvae that were at instars L3 & L4 during the previous cohort had hatched for the subsequent cohort. Consequently, each cohort contained a representative group of a colony’s worker production to identify infection dynamics over time. To investigate the impact of caffeine on infection prevalence and intensity in foragers, which are the route for inter-colony transmission of the parasite, every seven days 10% of the total colony population that were foraging were removed and screened for *N. bombi* infection via microscopic examination of the faeces. Each forager had its brood cohort and colony recorded. If an infection was identified a Neubauer improved haemocytometer was used to calculate the infection intensity (cells/μl). Finally, when the colony was terminated all remaining bees were removed and isolated in individual, labeled, 25ml collection tubes and workers were screened for *N. bombi* infection as described above. For each bee a record of sex, brood cohort and colony was taken, before they were sacrificed in a −80^°^C freezer. To ensure consistency across colonies with respect to termination, the following parameters were used to define the end of a colony. Colony endpoint was defined as either three weeks following the death of the original queen, three weeks following the eclosure of the first sexual caste or three weeks after the queen last laid a clutch of eggs. These guidelines ensured that all developing brood would reach eclosure by the endpoint, enabling a robust estimation of the complete reproductive output of a colony.

### Statistical analysis

All statistical analyses and graphical outputs were undertaken in R open-source programming language [82, 83]. Chi-squared (χ^2^) tests were used to compare the infection prevalence between treatments in both the therapeutic and prophylactic investigations. To analyse the therapeutic and prophylactic effect of caffeine on *N. bombi* infection intensity in newly eclosed workers, two separate linear mixed-effects models (LMM) were constructed. Models were constructed in the R package ‘lme4’ [84]. Both models were constructed under the following parameters. Infection intensity was used as a response variable, with treatment group as a categorical factor, in addition thorax width (mm) and faeces volume (μl) were included as covariates. Both models also incorporated colony-of-origin as a random effect. In addition, to analyse the impact of nectar caffeine on *N. bombi* epidemiology in *B. terrestris* colonies two LMM were created. The *N. bombi* prevalence model was constructed by having infection state (0/1) as a response variable, with phytochemical treatment as a categorical factor and number of inoculated brood, number of days since inoculation and brood cohort as covariates. Similarly, the *N. bombi* infection intensity model was constructed by having infection intensity (cells/μl) as a response variable with phytochemical treatment as a categorical factor and nuumber of inoculated brood, number of days since inoculation and brood cohort as covariates. In addition, this model also included cohort and treatment as an interaction. Both models also incorporated colony-of-origin and forager (0/1) as random effects. To analyse the prevalence and infection intensity of *N. bombi* in *B. terrestris* foragers two further LMM were constructed. Similar to above, the *N. bombi* prevalence model was constructed by having infection state (0/1) and the infection intensity model had cells/μl as response variables. For these models, phytochemical treatment was included as a categorical factor and number of inoculated brood, and brood cohort were covariates, with colony being included as a random factor. Finally, two LMM were constructed to investigate impact of caffeine treatment on colony fitness. To analyse colony size, worker production since inoculation was used as a response variable with treatment as a categorical factor and days since inoculation and number of brood inoculated as covariates. To analyse the production of reproductive castes, sexual production was used as a response variable with treatment as a categorical factor and number of brood inoculated, number of days since inoculation and number of workers at colony terminus as covariates. Again, both models included colony as a random effect. For infection intensity response models count data was log transformed and all models were validated in R by visually checking normality of residuals, and for overdispersion and collinearity of variables.

## Acknowledgements

We would like to thank Lucy Thursfield and Sue Baldwin for technical support and Harry Siviter for statistical advice. We would also like to thank Emorsgate seeds for allowing us to sample flowers on their land, and Windsor Great Park for allowing us to collect wild bumblebees. AJF would also like to thank Isla for enriching his life.

## Author contributions

AJF, PCS and MJFB devised and designed the experiment. IWF assisted with phytochemical identification and HK assisted with flower sampling and phytochemical identification. AJF was responsible for carrying out the experimental work and writing the manuscript. All authors provided feedback on the manuscript and agreed to its publication.

## Data availability

Data has been made open access and deposited onto FigShare under the title ‘Agri-environment scheme nectar chemistry negatively impacts bumblebee parasite epidemiology’ doi:10.6084/m9.figshare.13668767

## Supplementary files

**Supplementary table 1.**
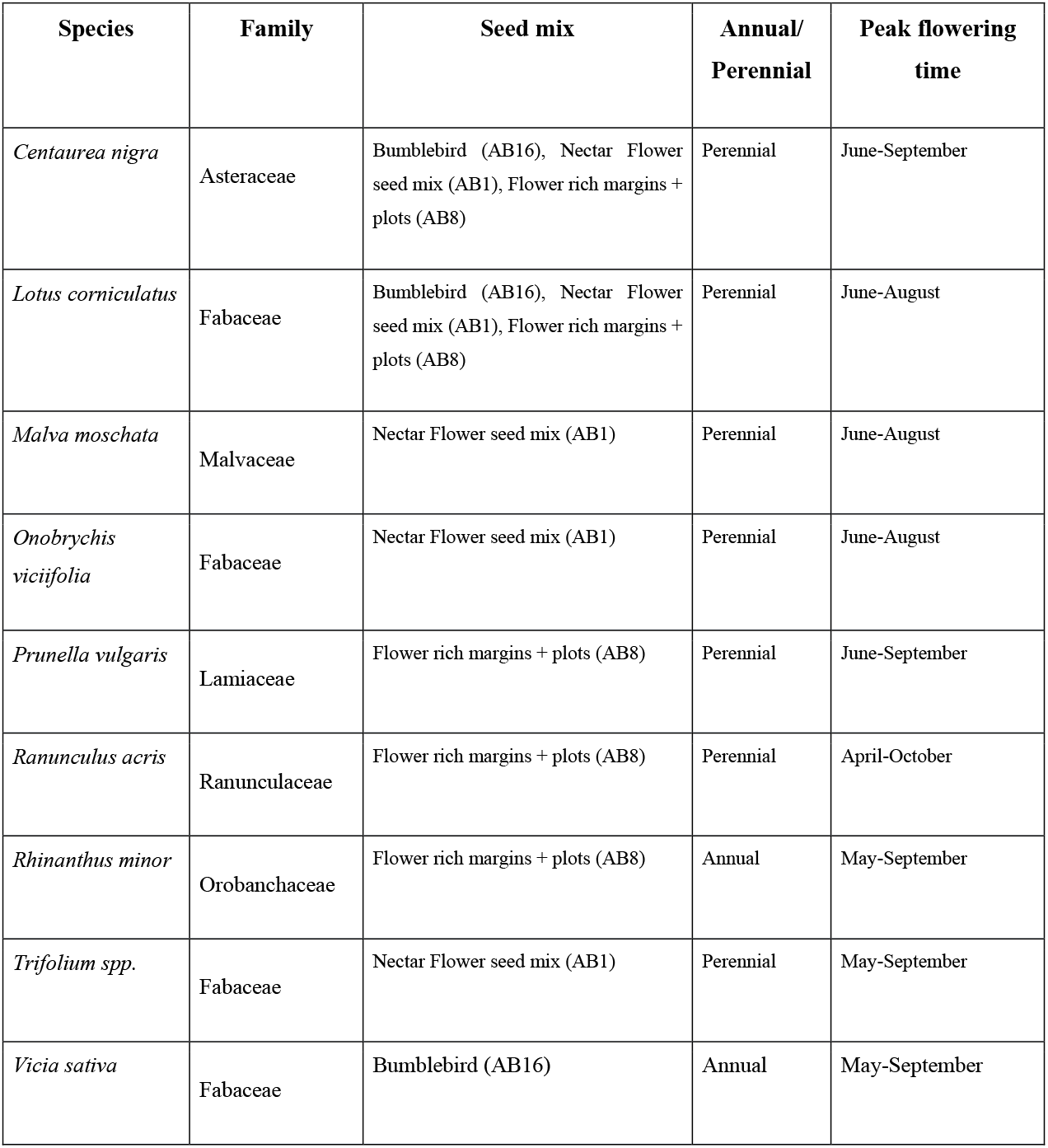
Flowers which are visited by bumblebees, from seed mixes, outlined by Natural England 2017, and that are recommended for use in UK based Agri-environment schemes. Other plants contained in the mixes, not listed or tested here, are selected for birds, hoverflies and other nectarivorous animals.

| Species | Family | Seed mix | Annual/<br>Perennial | Peak flowering<br>time |
| --- | --- | --- | --- | --- |
| *Centaurea nigra* | Asteraceae | Bumblebird (AB16), Nectar Flower seed mix (AB1), Flower rich margins + plots (AB8) | Perennial | June-September |
| *Lotus corniculatus* | Fabaceae | Bumblebird (AB16), Nectar Flower seed mix (AB1), Flower rich margins + plots (AB8) | Perennial | June-August |
| *Malva moschata* | Malvaceae | Nectar Flower seed mix (AB1) | Perennial | June-August |
| *Onobrychis viciifolia* | Fabaceae | Nectar Flower seed mix (AB1) | Perennial | June-August |
| *Prunella vulgaris* | Lamiaceae | Flower rich margins + plots (AB8) | Perennial | June-September |
| *Ranunculus acris* | Ranunculaceae | Flower rich margins + plots (AB8) | Perennial | April-October |
| *Rhinanthus minor* | Orobanchaceae | Flower rich margins + plots (AB8) | Annual | May-September |
| *Trifolium spp.* | Fabaceae | Nectar Flower seed mix (AB1) | Perennial | May-September |
| *Vicia sativa* | Fabaceae | Bumblebird (AB16) | Annual | May-September |

**Supplementary table 2.**
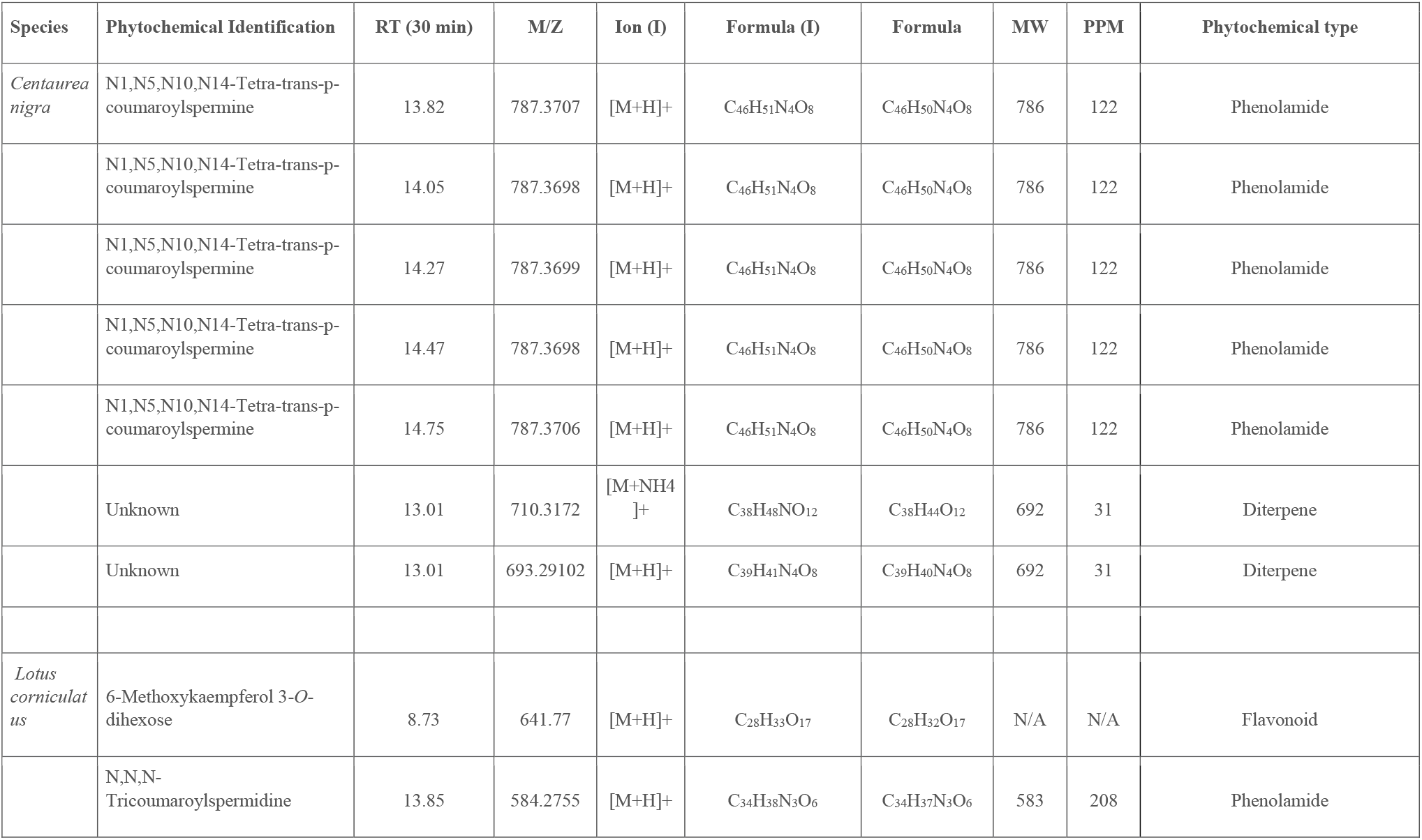
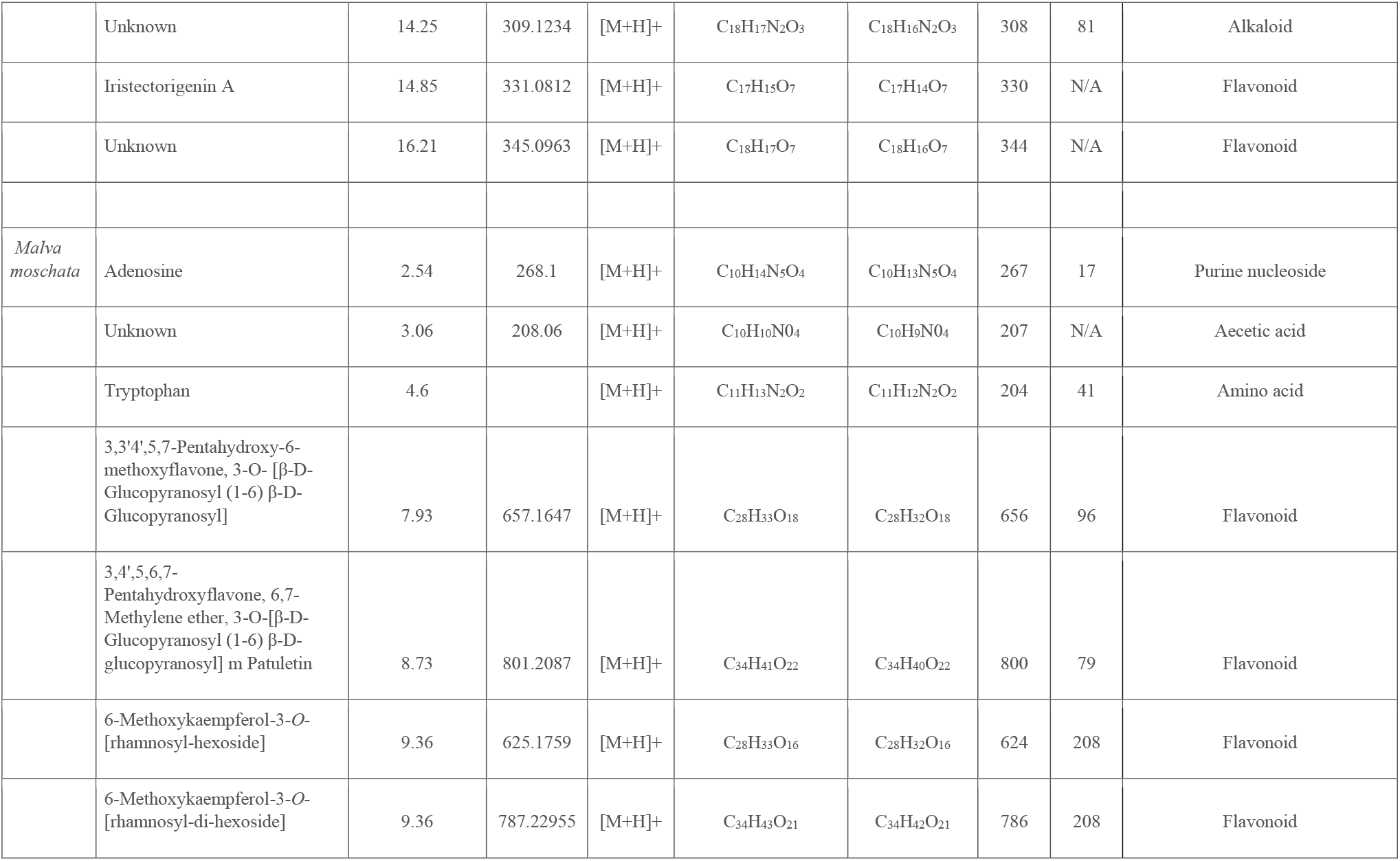
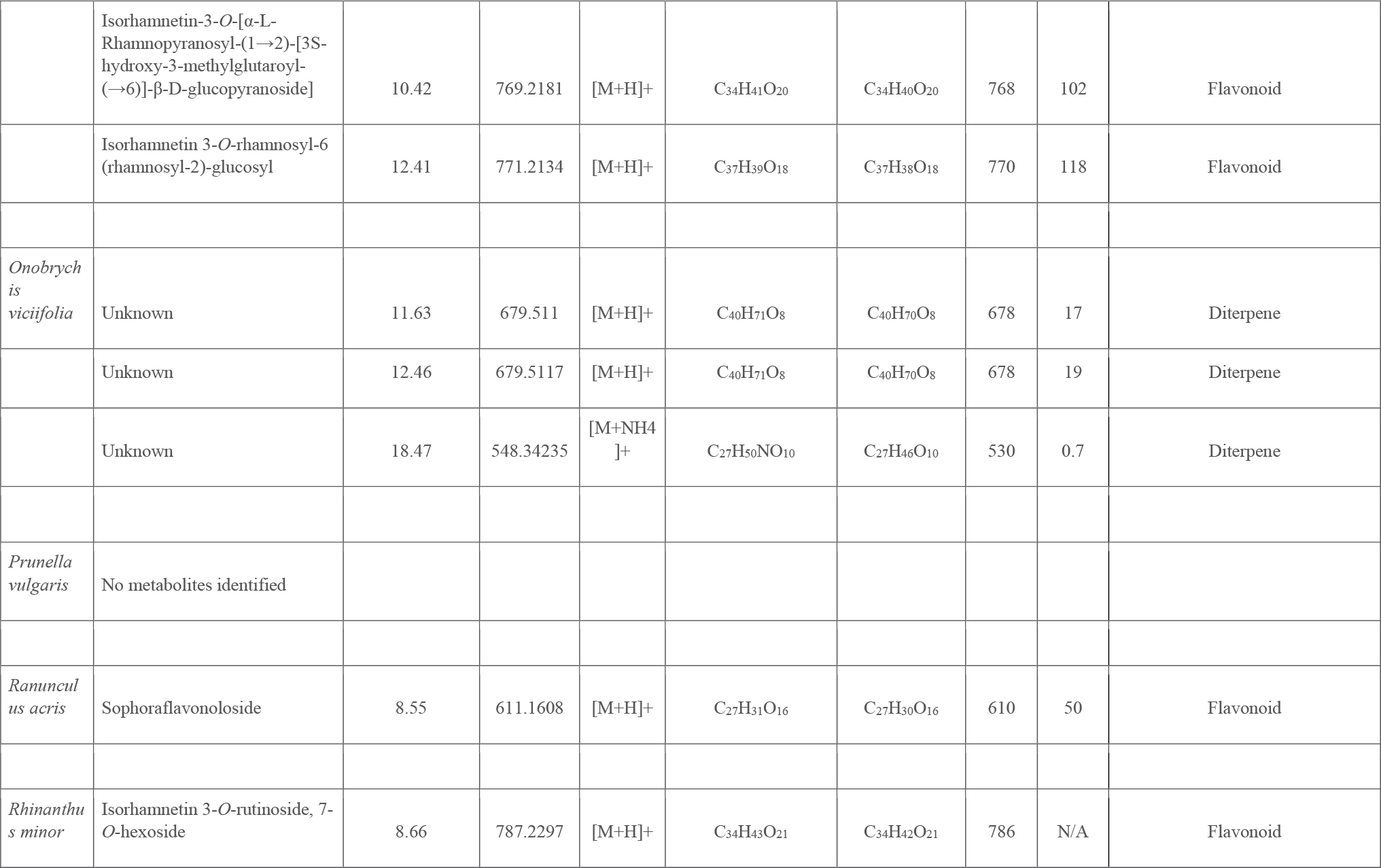
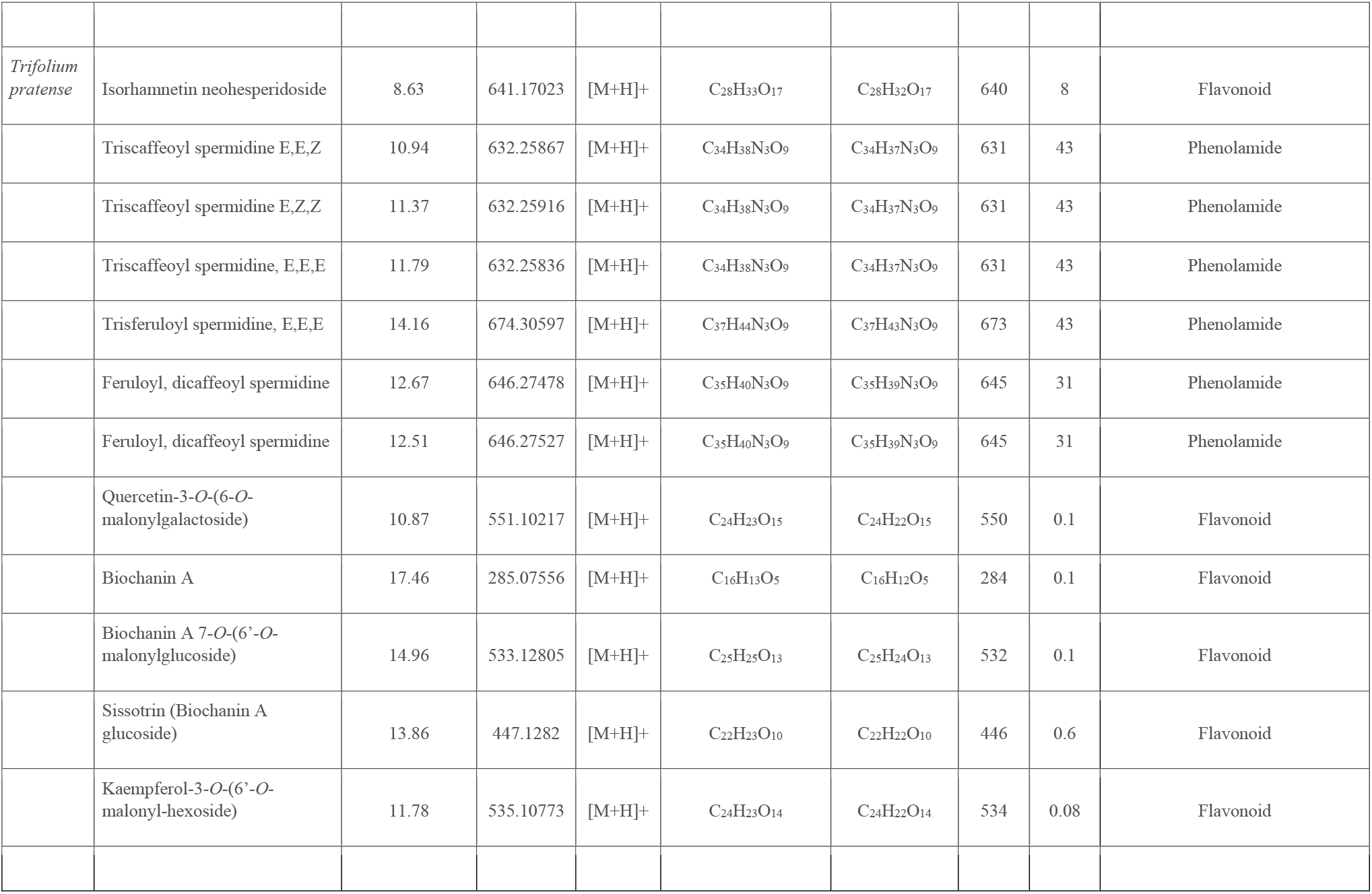
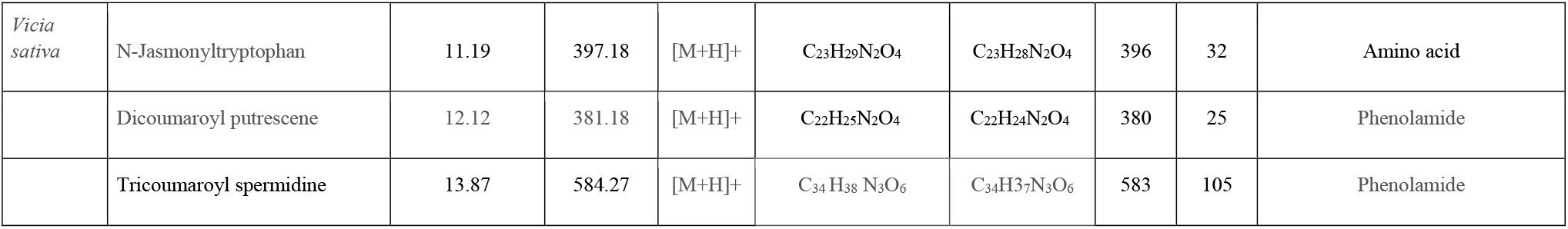
Phytochemicals recovered using LC-MS from the pollen of plants in UK agri-environment schemes.

**Supplementary table 3.**
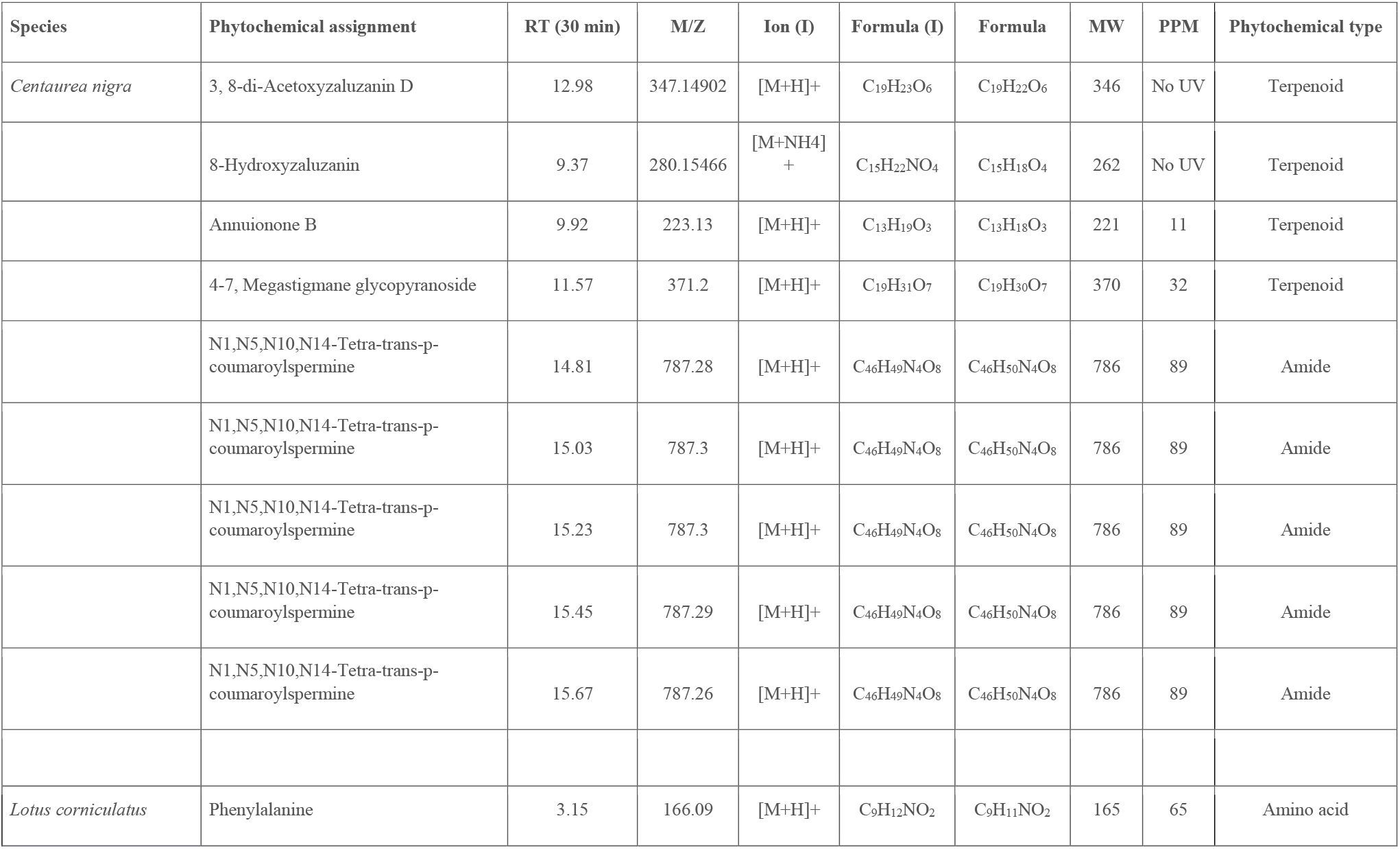
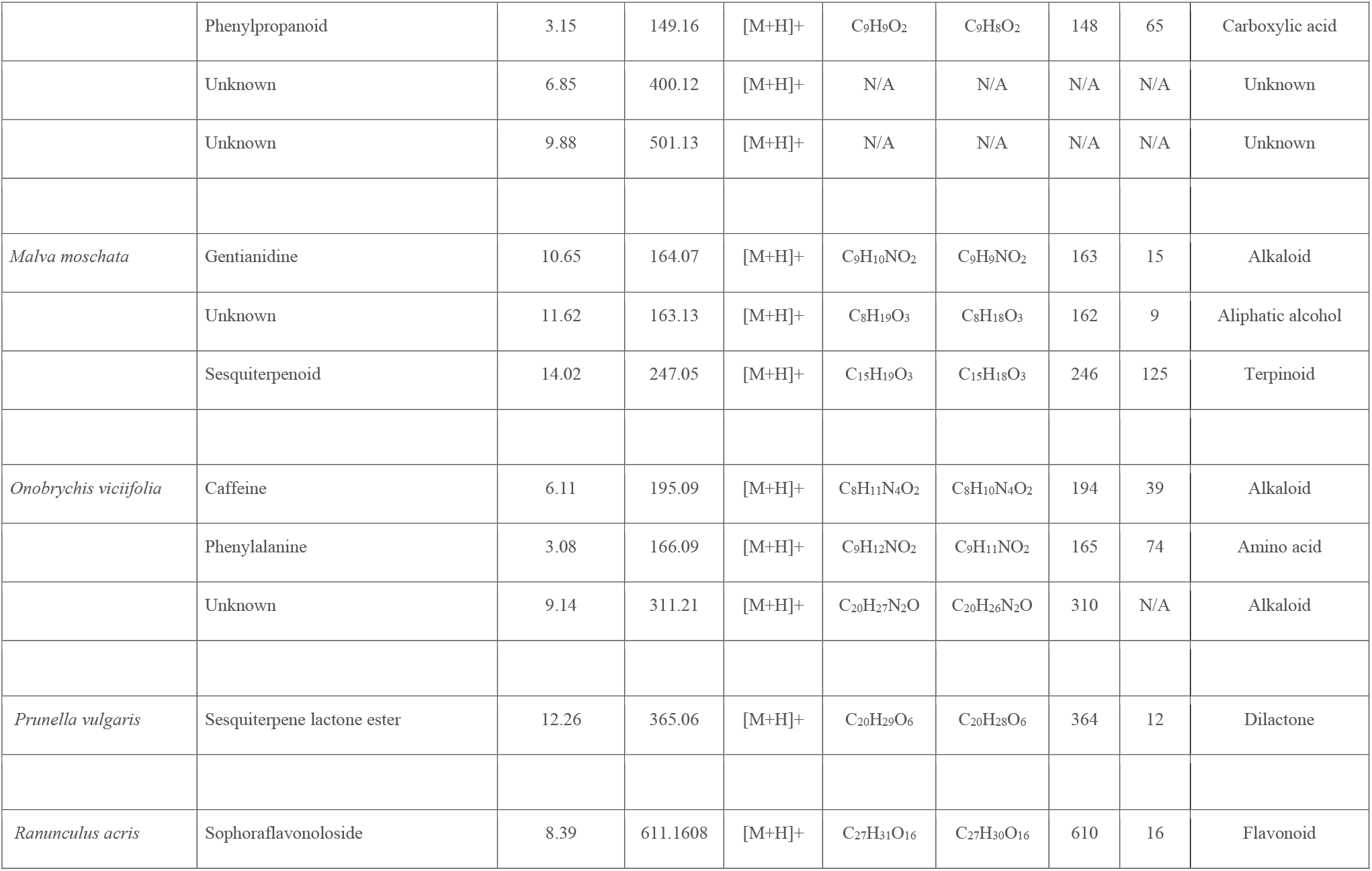
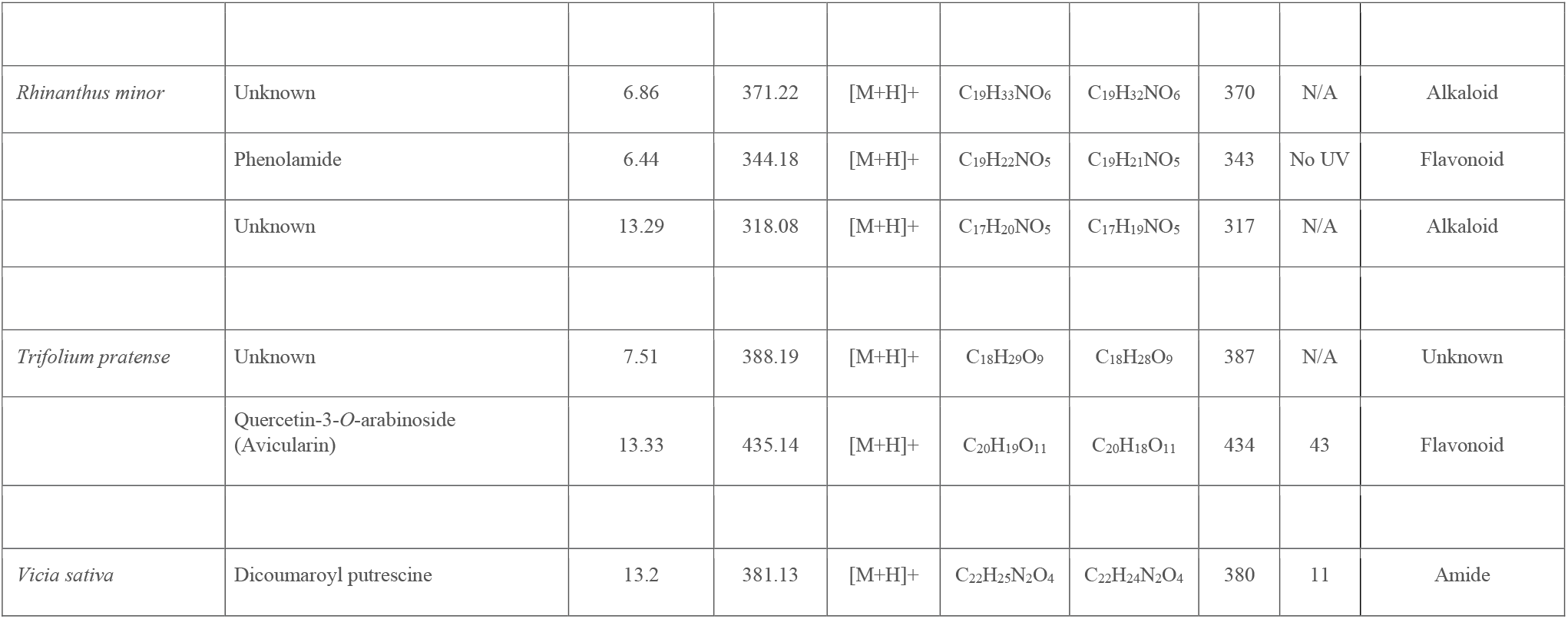
Phytochemicals recovered using LC-MS from the nectar of plants in UK based agri-environment schemes.

## References

[1] Barnosky, A.D., Matzke, N., Tomiya, S., Wogan, G.O.U., Swartz, B., Quental, T.B., Marshall, C., McGuire, J.L., Lindsey, E.L., Maguire, K.C., Mersey, B., Ferrer, E.A. (2011) Has the Earth’s sixth mass extinction already arrived? Nature. 471: 51–57.

[2] Berger, L., Speare, R., Daszak, P., Green, P.E., Cunningham, A.A., Goggin, C.L., Slocombe, R., Ragan, M.A., Hyatt, A.D., McDonald, K.R., Hines, H.B., Lips, K.R., Marantelli, G., Parkes, H. (1998) Chytridiomycosis causes amphibian mortality associated with population declines in the rain forests of Australia and Central America. Proceedings of the National Academy of Sciences of the United States of America. 95: 9031–9036.

[3] Daszak, P., Cunningham, A.A., Hyatt, A.D. (2000) Emerging infectious diseases of wildlife-Threats to biodiversity and human health. Science. 287: 1756.

[4] Jones, K.E., Patel, N.G., Leug, M.A., Storeygard, A., Balk, D., Gittleman, J.L., Daszak, P. (2008) Global trends in emerging infectious diseases. Nature. 451: 990–993.

[5] Leopardi, S., Blake, D., Puechmaille, S.J. (2011) White-Nose Syndrome fungus introduced from Europe to North America. Current Biology. 25: 217–219.

[6] Fisher, M.C., Garner, T.W.J. (2020) Chytrid fungi and global amphibian declines. Nature Reviews Microbiology. 18:332–343.

[7] Baude, D., Qi, X., Nielsen-Saines, K., Musso, D., Pomar, L., Favre, G. (2020) Real estimates of mortality following COVID-19 infection. The Lancet Infectious Diseases. 7: p773.

[8] Tilman, D., May, R.M., Lehman, C.L., Nowak, M.A. (1994) Habitat destruction and the extinction debt. Nature. 371: 65–66

[9] Donald, P.F., Green, R.E., Heath, M.F. (2001) Agricultural intensification and the collapse of Europe’s farmland bird populations. Proceedings of the Royal Society B. 268: 25–29.

[10] Travis, J.M.J. (2003) Climate change and habitat destruction: a deadly anthropogenic cocktail. Proceedings of the Royal Society B. 270: 467–473.

[11] Ollerton, J., Erenler, H., Edwards, M., Crockett, R. (2014) Pollinator declines. Extinctions of aculeate pollinators in Britain and the role of large-scale agricultural changes. Science. 346: 1360–1362.

[12] Senapathi, D., Carvalheiro, L.G., Biesmeijer, J.C., Dodson, C.A., Evans, R.L., McKerchar, M., Morton, R.D., Moss, E.D., Roberts, S.P.M., Kunin, W.E., Potts, S.G. (2015) The impact of over 80 years of land cover changes on bee and wasp pollinator communities in England. Proceedings of the Royal Society B. 282: 20150294.

[13] Powney, G.D., Carvell, C., Edwards, M., Morris, R.K.A., Roy, H.E., Woodcock, B.A., Isaac, N.J.B. (2019) Widespread losses of pollinating insects in Britain. Nature Communications. 10: 1018.

[14] Krebs, J.R., Wilson, J.D., Bradbury, R.B., Siriwardena, G.M. (1999) The second silent spring? Nature. 400: 611–612.

[15] Robinson, R.A., Sutherland, W.J. (2002) Post-war changes in arable farming and biodiversity in Great Britain. Journal of Applied Ecology. 39: 157–176.

[16] Reidsma, P., Tekelenburg, T., van der Berg, M., Alkemade, R. (2006) Impacts of land use change on biodiversity: An assessment of agricultural biodiversity in the European Union. Agriculture, Ecosystems and the Environment. 114: 86–102

[17] EU Common Agricultural Policy (2015). Brussels.

[18] Natural England (2017) Countryside Stewardship Manual version 27. Natural England.

[19] USDA Farm Service Agency (2016) Conservation Reserve Program. Farm Service Agency.

[20] Pywell, R.F., Warman, E.A., Hulmes, L., Hulmes, S., Nuttall, P., Sparks, T.H., Critchley, C.N.R., Sherwood, A. (2006) Effectiveness of new agri-environment schemes in providing foraging resources for bumblebees in intensively farmed landscapes. Biological Conservation. 129:192–206.

[21] Carvell, C., Meek, W.R., Pywell, R.F., Goulson, D., Nowakowski, M. (2007) Comparing the efficacy of agri environment schemes to enhance bumble bee abundance and diversity on arable field margins. Journal of Applied Ecology. 44: 29–40.

[22] Wood, T.J., Holland, J.M., Hughes, W.O.H., Goulson, D. (2015) Targeted agri-environment schemes significantly improve the population size of common farmland bumblebee species. Molecular Ecology. 24: 1668–1680.

[23] Carvell, C., Bourke, A.F.G., Dreier, S., Freeman, S.N., Hulmes, S., Jordan, W.C., Redhead, J.W., Sumner, S., Wang, J., Heard, M.S. (2017) Bumblebee family lineage survival is enhanced in high quality landscapes. Nature. 543: 547–548.

[24] Cameron, S.A., Lozier, J.D., Strange, J.P., Koch, JB., Cordes, N., Solter, L.F., Griswold, T.L. (2011) Patterns of widespread decline in North American bumble bees. Proceedings of the National Academy of Sciences of the United States of America. 108: 662–667.

[25] Fürst, M.A., McMahon, D.P., Osborne, J.L., Paxton, R.J., Brown, M.J.F. (2014) Disease associations between honeybees and bumblebees as a threat to wild pollinators. Nature. 506: 364.

[26] McArt, S.H., Koch, H., Irwin, R.E., Adler, L.S. (2014) Arranging the bouquet of disease: floral traits and the transmission of plant and animal pathogens. Ecology Letters. 17: 624–636.

[27] Bailes, E., Bagit, J., Coltman, J., Fountain, M.T., Wilfert, L., Brown, M.J.F. (2020) Host density drives viral, but not trypanosome, transmission in a key pollinator. Proceedings of the Royal Society B. 287:20191969.

[28] Baker, H.G., Baker, I. (1975) Studies of nectar constitution and pollinator-plant coevolution. Coevolution of plants and animals. University of Texas Press.

[29] Adler, L.S. (2000) The ecological significance of toxic nectar. Oikos. 91: 409–420.

[30] Stevenson, P.C., Nicolson, S.W., Wright, G.A. (2017) Plant secondary metabolites in nectar: Impacts on pollinators and ecological functions. Functional Ecology. 31: 65–75.

[31] Cowan, M.M. (1999) Plant products as antimicrobial agents. Clinical Microbiology Reviews. 12: 564–582.

[32] Manson, J.S., Otterslatter, M.C., Thomson, J.D. (2010) Consumption of a nectar alkaloid reduces pathogen load in bumble bees. Oecologia. 162: 81–89.

[33] Richardson, L.L., Adler, L.S., Leonard, A.S., Andicoechea, J., Regan, K.H., Anthony, W. E., Manson, J.S., Irwin, R.E. (2015) Secondary metabolites in floral nectar reduce parasite infections in bumblebees. Proceedings of the Royal Society B. 282: 20142471.

[34] Giacomini, J.J., Leslie, J., Tarpy, D.R., Palmer-Young, E.C., Irwin, R.E., Adler, L.S. (2018) Medicinal value of sunflower pollen against bee pathogens. Scientific reports. 8: 14394.

[35] Koch, H., Woodward, J., Langat, M.K., Brown, M.J.F., Stevenson, P.C. (2019) Flagellum removal by a nectar metabolite inhibits infectivity of a bumblebee parasite. Current Biology. 29: 3494–3500.

[36] Folly, A.J., Stevenson, P.C., Brown, M.J.F. (2020) Age-related pharmacodynamics in a bumblebee microsporidian system mirror similar patterns in vertebrates. Journal of Experimental Biology. 223: jeb217828.

[37] Breeze, T.D., Bailey, A.P., Balcombe, K.G., Potts, S.G. (2011) Pollination services in the UK: How important are honeybees. Agriculture, Ecosystems and Environment. 142: 137–143.

[38] Kleijn, D., Winfree, R., Bartomeus, I., Carualheiro, L.G., Henry, M., Isaacs, R., Klein, A.M., Kremen, C., M’Gonigle, L.K., Rander, R., Ricketts, T.H., Williams, N.H., Adamson, N.L., Ascher, J.S., Báldi, A., Batáry, P., Benjamin, F., Beismeijer, J., Blitzer, E.J., Bommarco, R., Brand, M.R., Bretagnolle, V., Button, L., Cariveau, D.P., Chifflet, R., Colville, J.F., Danforth, B.N., Elle, E., Garratt, M.P.D., Herzog, F., Holzschuh, A., Howlett, B.G., Jauker, F., Jha, S., Knop, E., Krewenka, K.M., Le Féon, V., Mandelik, Y., May, E.A., Park, M.G., Pisanty, G., Reemer, M., Sardiñas, H.S., Scheper, J., Sciligo, A.R., Smith, H.G., Steffan-Dewenter, I., Thorp, R., Tscharntke, T., Verhulst, J., Vianna, B.F., Vaissière, B.E., Veldtman, R., Ward, K.L., Westphal, C., Potts, S.G. (2015) Delivery of crop pollination services is an insufficient argument for wild pollinator conservation. Nature Communications. 6: 7414.

[39] Woodcock, B.A., Garratt, M.P.D., Powney, G.D., Shaw, R.F., Osborne, J.L., Soroka, J., Lindström, S.A.M., Stanley, D., Ouvard, P., Edwards, M.E., Jauker, F., McCracken, M.E., Zou, Y., Potts, S.G., Rundlöf, Noriega, J.A., Greenop, A., Smith, H.G., Bommarco, R., van der Werf, W., Stout, J.C., Steffan-Dewenter, I., Morandin, L., Bullock, J.M., Pywell, R.F. (2019) Meta-analysis reveals that pollinator functional diversity and abundance enhance crop pollination and yield. Nature Communications. 10:1481.

[40] Vanbergen, A.J., the Insect Pollinators Initative. (2013) Threats to an ecosystem service: pressures on pollinators. Frontiers in Ecology and the Environment. 11: 251–259.

[41] Brown, M.J.F. (2017) Microsporidia: An emerging threat to bumblebees? Trends in Parasitology. 33: 754–762.

[42] McMahon, D.P., Fürst, M.A., Caspar, J., Theodorou, P., Brown, M.J.F., Paxton, R.J. (2015) A sting in the spit: widespread cross-infection of multiple RNA viruses across wild and managed bees. Journal of Animal Ecology. 84: 615–624.

[43] Schmid-Hempel, R., Eckhardt, M., Goulson, D., Heinzmann, D., Lange, C., Plischuck, S., Escudero, L.R., Salathé, R., Scriven, J.J., Schmid-Hempel, P. (2014) The invasion of South America by imported bumblebees and associated parasites. Journal of Animal Ecology. 83: 823–837.

[44] Fantham, H.B., Porter, A. (1914) The morphology, biology and economic importance of *Nosema bombi* n. sp. parasitic in various humble bees (*Bombus sp.*). Annals of Tropical Medicine and Parasitology. 8: 623–638.

[45] Cameron, S.A., Lim, H.C., Lozier, J.D., Duennes, M.A., Thorp, R. (2016) Test of the invasive pathogen hypothesis of bumblebee decline in North America. Proceedings of the National Academy of Sciences of the United States of America. 113: 4386–4391.

[46] Munday, Z., Brown, M.J.F. (2018) Bring out your dead: quantifying corpse removal in *Bombus terrestris*, an annual eusocial insect. Animal Behaviour. 138: 51–57.

[47] Cremer, S.M Armitage, S.A.O., Schmid-Hempel, P. (2007) Social Immunity. Current Biology. 17: 693–702.

[48] Huffman, M.A. (2001) Self-medicative behaviour in the African great apes: An evolutionary perspective into the origins of human traditional medicine. BioScience. 51: 651–661.

[49] de Roode, J.C., Pedersen, A.B., Hunter, M.D., Altizer, S. (2007) Host plant species affects virulence in monarch butterfly parasites. Journal of Animal Ecology. 77: 120–126.

[50] de Roode, J.C., Lefèvre, T., Hunter, M.D. (2013) Self medication in animals. Science. 340: 150.

[51] Huffman, M.A., Seifu, M. (1989) Observations of the illness and consumption of a possible medicinal plant *Vernonia amygdalina* (Del), by Chimpanzees in the Mahale Mountains National Park, Tanzania. Primates. 30: 51–63.

[52] Huffman, M.A., Gotoh, S., Izutsu, D., Koshimizu, K., Kalunde, M.S. (1993) Further observations on the use of *Vernonia amygdalina* by wild Chimpanzees, its possible effect on parasite load, and its phytochemistry. African Study Monographs. 14: 227–240.

[53] Sternberg, E.D., Lefévre, T., li, J., Fernandez de Castillejo, C.L., Li, H., Hunter, M.D., de Roode, J.C. (2012) Food plant-derived disease tolerance and resistance in a natural butterfly-plant-parasite interactions. Evolution. 66: 3367–3376.

[54] Schmid-Hempel, P. (1998) Parasites in social insects. Princeton University Press, Princeton. New Jersey.

[55] Food and Agriculture Organisation of the United Nations (1998) FAOSTAT Statistics Database. Rome.

[56] Wright, G.A., Baker, D.D., Palmer, M.J., Stabler, D., Mustard, J.A., Power, E.F., Borland, A.M., Stevenson, P.C. (2013) Caffeine in floral nectar enhances a pollinators memory of reward. Science. 339: 1202.

[57] Huang, R., Donnell, A.J., Barboline, J.J., Barkman, T.J. (2016) Convergent evolution of caffeine in plants by co-option of exapted ancestral enzymes. Proceedings of the National Academy of Sciences of the United States of America. 113: 10613–10618.

[58] Egan, P.A., Stevenson, P.C., Tiedeken, E.J., Wright, G.A., Boylan, F., Stout, J.C. (2016) Plant toxin levels in nectar vary spatially across native and introduced populations. Journal of Ecology. 104:1106–1115.

[59] Raj, C.V., Dhala, S. (1965) effect of naturally occurring xanthines on bacteria. I. antimicrobial action and potentiating effect on antibiotic spectra. Applied Microbiology. 13: 432–436.

[60] Hsieh, E.M., Berenbaum, M.R., Dolezal, A.G. (2020) Ameliorative effects of phytochemical ingestion on viral infection in Honey Bees. Insects. 11: 698.

[61] Otti, O., Schmid-Hempel, P. (2008) A field experiment on the effect of *Nosema bombi* in colonies of the bumblebee *Bombus terrestris*. Ecological Entomology. 33: 577–582.

[62] Shykoff, J., Schmid-Hempel, P. (1991) Incidence and effects of four parasites in natural populations of bumblebees in Switzerland. Apidologie. 22: 117–125.

[63] Durrer, S., Schmid-Hempel, P. (1994) Shared use of flowers leads to horizontal pathogen transmission. Proceedings of the Royal Society B. 258: 299–302.

[64] Ruiz-González, M.X., Bryden, J., Moret, Y., Reber-Funk, C., Schmid-Hempel, P., Brown, M.J.F. (2012) Dynamic transmission, host quality and population structure in a multihost parasite of bumblebees. Evolution. 66: 3053–3066.

[65] Free, J.B. (1955) The division of labour within bumblebee colonies. Insectes Sociaux. 2: 195–212.

[66] Folly, A.J., Koch, H., Stevenson, P.C., Brown, M.J.F. (2017) Larvae act as a transient transmission hub for the prevalent bumblebee parasite *Crithidia bombi*. Journal of Invertebrate Pathology. 148: 81–85.

[67] Rutrecht, S. T., Klee, J., Brown, M. J. F. (2007). Horizontal transmission success of *Nosema bombi* to its adult bumble bee hosts: effects of dosage, spore source and host age. Parasitology 134(12), 1719–1726.

[68] Otti, O., Schmid-Hempel, P. (2007) *Nosema bombi:* A pollinator parasite with detrimental fitness effects. Journal of Invertebrate Pathology. 96: 118–124.

[69] Arnold, S.E.J., Idrovo, E.P., Lomas Arias, L.J., Belmain, S.T., Stevenson, P.C. (2014) Herbivore defense compounds occur in pollen and reduce bumblebee colony fitness. Journal of Chemical Ecology. 40: 878–881.

[70] Thomson, J.D., Draguleasen, M.A., Tan, M.G. (2015) Flowers with caffeinated nectar receive more pollination. Arthropod-Plant Interactions. 9: 1–7.

[71] Couvillon, M.J., Toufailia, H.A., Butterfield, T.M., Schrell, F., Ratnieks, F.L.W., Schürch, R. (2015) Caffeinated forage tricks honeybees into increasing foraging and recruitment behaviours. Current Biology. 25: 1–4.

[72] Pywell, R.F., Meek, W.R., Hulmes, L., Hulmes, S., James, K.L., Nowakowski, M., Carvell, C. (2011) Management to enhance pollen and nectar resources for bumblebees and butterflies within intensely farmed landscapes. Journal of Insect Conservation. 15: 853–864.

[73] Dicks, L.V., Baude, M., Roberts, S.P.M., Phillips, J., Green, M., Carvell, C. (2015) How much flower-rich habitat is enough for wild pollinators? Answering a key policy question with incomplete knowledge. Ecological Entomology. 40: 22–35.

[74] Corbet, S.A. (2003) Nectar sugar content: estimating standing crop and secretion rate in the field. Apidologie. 34: 1–10.

[75] Morrant, D.S., Schumann, R., Petit, S. (2009) Field methods for sampling and storing nectar from flowers with low nectar volumes. Annals of Botany. 103: 533–542.

[76] Palmer-Young, E.C., Farrell, I.W., Adler, L.S., Milano, N.J., Egan, P.A., Junker, R.R., Irwin, R.E., Stevenson, P.C. (2018) Chemistry of floral rewards: intra-and interspecific variability of nectar and pollen secondary metabolites across taxa. Ecological Monographs. 89:doi.org/10.1002/ecm.1335.

[77] Cook, D., Manson, J.S., Gardner, D.R., Welch, K.D., Irwin, R.E. (2013) Norditerpine alkaloid concentrations in tissues and floral rewards of larkspurs and impacts on pollinators. Biochemical Systematics and Ecology. 48: 123–131.

[78] Islam, M.K., Sohrab, M., Jabbar, A. (2011) Caffeine and P-Anisaldehyde from the fruits of Enterolobium saman prain. International Journal of Pharmaceutical Sciences and Research. 3: 168–170.

[79] Rutrecht, S.T., Brown, M.J.F. (2008) Within colony dynamics of *Nosema bombi* infections: disease establishment, epidemiology and potential vertical transmission. Apidologie. 39: 504–514.

[80] Erler, S., Lommatzsch, S., Lattorff, M.G. (2012) Comparative analysis of detection limits and specificity of molecular diagnostic markers for three pathogens (Microsporidia, *Nosema spp.*)in the key pollinators *Apis mellifera* and *Bombus terrestris*. Parasitology Research. 110: 1403–1410.

[81] Logan, A., Ruiz-González, M.X., Brown, M.J.F. (2005) The impact of host starvation on parasite development and population dynamics in an intestinal trypanosome parasite of bumblebees. Parasitology. 130: 637–642.

[82] R Core Team (2020) R: A language and environment for statistical computing. R Foundation for Statistical Computing, Vienna, Austria.

[83] Wickham, H. (2009) ggplot 2: Elegant graphics for data analysis. Springer-Verlag. New York.

[84] Bates, D., Maechler, M., Bolker, B., Walker, S. (2015) Fitting linear mixed effects models using Lme4. Journal of Statistical Software. 67: 1–48.

